# *In vitro* reconstitution reveals requirement of Nek2a and cyclin A2 for Wapl-dependent removal of cohesin from prophase chromatin

**DOI:** 10.1101/2024.03.12.584565

**Authors:** Susanne Hellmuth, Olaf Stemmann

## Abstract

Sister chromatid cohesion is mediated by the cohesin complex. In mitotic prophase cohesin is removed from chromosome arms in a Wapl- and phosphorylation-dependent manner. Sgo1-PP2A protects pericentromeric cohesion by dephosphorylation of cohesin and its associated Wapl antagonist sororin. However, Sgo1-PP2A relocates to inner kinetochores well before sister chromatids are separated by separase, leaving pericentromeric regions unprotected. Why deprotected cohesin is not removed by Wapl remains enigmatic. By reconstituting Wapl-dependent cohesin removal from chromatin *in vitro*, we discovered a requirement for Nek2a and Cdk1/2-cyclin A2. These kinases phosphorylate cohesin-bound Pds5b, thereby converting it from a sororin- to a Wapl- interactor. Replacement of endogenous Pds5b by a phosphorylation mimetic variant causes premature sister chromatid separation (PCS). Conversely, phosphorylation-resistant Pds5b impairs chromosome arm separation in prometaphase-arrested cells and suppresses PCS in the absence of Sgo1. Early mitotic degradation of Nek2a and cyclin A2 may therefore explain why only separase, but not Wapl, can trigger anaphase.

## Introduction

The ring-shaped cohesin complex mediates sister chromatid cohesion by topologically embracing two DNA molecules in its center (Gruber, Haering et al., 2003, Haering, Farcas et al., 2008, Ochs, Green et al., 2024). It also structures chromatin by DNA loop extrusion - but this is not further discussed here. Cohesin consists of four subunits, which in mitotically dividing human cells are the two structural maintenance of chromosomes (SMC) proteins Smc1A and Smc3, the kleisin Rad21 and one of two related stromal antigens, Stag1/SA1 or Stag2/SA2 (Losada, Hirano et al., 1998, Losada, Yokochi et al., 2000). Smc1 and Smc3 each have an ABC-like ATPase head domain connected by an antiparallel, highly elongated coiled-coil to a central hinge domain that mediates Smc1-Smc3 heterodimerization (Anderson, Losada et al., 2002, Haering, Lowe et al., 2002). Rad21 interacts via its N-terminal helical domain (NHD) with the head-proximal coiled coil of Smc3 (the ‘neck’) and via its C-terminal winged helix domain (WHD) with the Smc1 head, thereby forming a tripartite SMC-kleisin (S-K) ring (Gligoris 2014; Haering, 2004). Engagement of the SMC heads by sandwiching of two ATP molecules between them creates smaller S and K rings, whereas their disengagement upon ATP hydrolysis reforms the large S-K ring (Shi, Gao et al., 2020). The NHD and WHD of Rad21 are connected by a flexible linker that in its C-terminal part is associated with Stag1 or -2 and in its N-terminal part with the more loosely bound NIPBL, Pds5a or Pds5b (Hara, Zheng et al., 2014, Kikuchi, Borek et al., 2016, Lee, Roig et al., 2016, Ouyang, Zheng et al., 2016). Because Stag/SA, NIPBL and Pds5 all share extensive HEAT repeats, they are commonly referred to as HAWKs (HEAT repeat proteins associated with kleisins) (Wells, Gligoris et al., 2017).

In human cells, cohesin is loaded in a NIPBL- and ATP-dependent manner onto one- chromatid chromosomes (Elbatsh, Haarhuis et al., 2016, Watrin, Schleiffer et al., 2006). This first occurs in telophase, but this association remains dynamic throughout G1 phase (Darwiche, Freeman et al., 1999, Gerlich, Koch et al., 2006). The co-replicative establishment of sister chromatid cohesion requires entrapment of a second DNA and is coupled with cohesion-stabilizing acetylation of the Smc3 head by Esco1/2 (Hou & Zou, 2005, Murayama, Samora et al., 2018). After exchange of NIPBL for Pds5a/b, the recruitment of sororin to acetylated cohesin competitively inhibits the binding of the anti-cohesive factor Wapl to Pds5 (Nishiyama, Ladurner et al., 2010). In addition, sororin is predicted to bind to the Rad21-Smc3 junction, which may contribute to its cohesion-protective function (see below) (Nasmyth, Lee et al., 2023). Cohesion allows mitotic spindle forces to be sensed and resisted, which facilitates proper amphitelic attachment of sister kinetochores. While this is critical for error-free chromosome segregation, all cohesive cohesin must be removed from chromatin before anaphase can begin. When metazoan cells enter mitosis, cohesin at chromosome arms is displaced by action of the so-called prophase pathway (Darwiche et al., 1999, Losada et al., 1998, Waizenegger, Hauf et al., 2000). This involves phosphorylation-dependent inactivation of sororin, which is then replaced by Wapl as a binding partner of Pds5 (Dreier, Bekier et al., 2011, Liu, Jia et al., 2013a, Nishiyama, Sykora et al., 2013). Rad21 then dissociates from Smc3 and DNA exits the ring through this opened exit gate (Buheitel & Stemmann, 2013, Chan, Roig et al., 2012, Eichinger, Kurze et al., 2013, Huis in ‘t Veld, Herzog et al., 2014). Exactly how Wapl unloads cohesin is still unclear, but it may act indirectly by sequestering Rad21’s NTD after its detachment from Smc3’s neck due to ATP-driven engagement of the SMC heads (Elbatsh et al., 2016, Muir, Li et al., 2020, Nasmyth et al., 2023). (Peri)centromeric cohesion is resistant to the Wapl-dependent release in mitotic prophase. This is because the cohesion protector shugoshin 1 (Sgo1) binds to pericentromeric cohesin upon its phosphorylation by Cdk1-cyclin B1 (Liu et al., 2013a, McGuinness, Hirota et al., 2005, Tang, Sun et al., 2004). Sgo1, in turn, recruits protein phosphatase 2A (PP2A), which maintains sororin (and cohesin) in a dephosphorylated state (Kitajima, Sakuno et al., 2006, Riedel, Katis et al., 2006, Tang, Shu et al., 2006). In addition, Sgo1 binds to the SA2-Rad21 contact site, thereby preventing the simultaneous association of Wapl with SA2 (Garcia-Nieto, Patel et al., 2023, Hara et al., 2014). Pericentromeric cohesion is dissolved only at the end of metaphase, when a giant protease, separase, is activated and cleaves Rad21 (Hauf, Waizenegger et al., 2001, Uhlmann, Wernic et al., 2000). This proteolytic opening of residual cohesin triggers sister chromatid separation and marks the onset of anaphase.

The prophase pathway is important for error-free chromosome segregation and the prevention of aneuploidies (Haarhuis, Elbatsh et al., 2013, Tedeschi, Wutz et al., 2013). Next to Wapl, this proteolysis-independent opening of cohesin rings at chromosome arms requires mitotic phosphorylations. Cdk1-cyclin B1, Plk1 and aurora B have been reported as the relevant kinases, while sororin and SA2 have been identifed as important substrates of phosphorylation (Dreier et al., 2011, Gimenez-Abian, Sumara et al., 2004, Hauf, Roitinger et al., 2005, Lenart, Petronczki et al., 2007, Liu et al., 2013a, Losada, Hirano et al., 2002, Nishiyama et al., 2013, Sumara, Vorlaufer et al., 2002). However, several issues remain. The prophase pathway has never been reconstituted in a purified system, thus leaving unanswered whether the three kinases and their two targets are necessary and sufficient for Wapl-driven release activity. Furthermore, amphitelic attachment of chromosomes to spindle microtubules coincides with relocalization of Sgo1-PP2A from pericentromeres to inner kinetochores, which later ensures unopposed cohesin cleavage by separase by removing a steric hindrance and allowing proteolysis-promoting phosphorylation of Rad21 (Hauf et al., 2005, Lee, Kitajima et al., 2008, Liu, Rankin et al., 2013b). This deprotection of cohesin and associated sororin occurs in prometaphase to metaphase, i.e. well before anaphase onset and at a time when Wapl is present and the above kinases are fully active. Why Wapl-induced premature sister chromatid separation (PCS) does not occur in a normal mitosis remains a mystery.

Here we reconstitute the human prophase pathway using purified recombinant proteins. We find that the combined action of the kinases Nek2a, Cdk1/2-cyclin A2 and aurora B are necessary and sufficient for Wapl-dependent release of cohesin and sororin from immobilized, high-salt washed G2 phase chromatin. Nek2a and Cdk1/2-cyclin A2 phosphorylate Pds5b, thereby converting it from a sororin- to a Wapl binder. Consistently, a Pds5b variant resistant to phosphorylation by these two kinases results in the retention of cohesin on chromosome arms of prometaphase-arrested cells and suppresses premature sister chromatid separation (PCS) in absence of Sgo1. Conversely, a Pds5b variant that mimics constitutive phosphorylation by Nek2a and Cdk1/2-cyclin A2 causes PCS. Taken together, our study reveals Nek2a and Cdk1/2-cyclin A2 as two critical factors of prophase pathway signaling and Pds5b as an important target of these kinases. Given the early mitotic degradation of Nek2a and cyclin A2 before deprotection of pericentromeric cohesin, our study also explains why anaphase is not triggered by Wapl but only by separase.

## Results

### Nek2a and Cdk1/2-cyclin A2 kinases are necessary for Wapl-dependent unloading of cohesin from isolated chromatin

To reconstitute the prophase pathway form purified components, we isolated recombinant Wapl, Cdk1-cyclin B1, Plk1 and aurora B-INCENP. Wapl readily bound to Pds5 (see below) and the kinases were active towards model substrates (Figure EV1A). However, these four components were not sufficient to displace cohesin and/or sororin from isolated G2- chromatin (Figure EV1B, lanes 1-3). Although pericentromeric cohesin is stripped from Sgo1- PP2A well before anaphase onset, Wapl cannot (and must not) trigger sister chromatid separation, at least in a healthy cell. To explain this conundrum, we hypothesized that the prophase pathway, as its name suggests, is no longer active in metaphase due to absence of required components. Nek2a and cyclin A2 are characterized by their early mitotic degradation and seemed to be candidates for the proposed, limiting factors (den Elzen & Pines, 2001, Geley, Kramer et al., 2001, Hames, Wattam et al., 2001). Therefore, we generated active Nek2a and Cdk1/2-cyclin A2 kinases (Figure EV1A) and included them in our release activity assays. Indeed, the addition of either kinase to the reconstitution reactions caused some cohesin and sororin, but not histone H2A, to be displaced from chromatin (Figure EV1B, lanes 4 + 5). Simultaneous addition of both kinases increased the release only slightly (Figure EV1B, lane 6). The reconstitution assay also allowed us to assess the necessity of Plk1, aurora B-INCENP and Cdk1-cyclin B1 for the displacement of cohesin from chromatin *in vitro*. Surprisingly, in the presence of Wapl, Nek2a and Cdk1/2-cyclin A2, aurora B-INCENP was necessary and sufficient for the release of cohesin and sororin from chromatin, whereas Cdk1-cyclin B1 and Plk1 were dispensable (Figure EV1C).

### Reconstitution of the prophase pathway using immobilized chromatin as substrate

To further improve our *in vitro* reconstitution system, we wanted to generate a biochemically more defined chromatin substrate. To this end, we used DNA-mediated chromatin pull-down (Dm-ChP) technology (Aranda, Alcaine-Colet et al., 2019, Kliszczak, Rainey et al., 2011). HeLaK cells pre-synchronized at the G1-S transition were released to undergo replication in presence of the alkine-containing thymidine analog F-ara-EdU (Neef & Luedtke, 2011). Following G2-arrest, cell lysis, and isolation of nuclei, the DNA was covalently labeled with azide-carrying biotin by a mild ‘click’ reaction and then immobilized on streptavidin sepharose (Figure 1A). The corresponding beads carried native chromatin as judged by their F-ara-EdU-dependent binding of histone H2A, cohesin, Pds5b, and sororin (Figure 1B, lanes 1-5). While histone H2A was removed by high salt, cohesin together with Pds5b and sororin were largely retained (Figure 1B, lane 6). Conversely and consistent with a topological entrapment of tethered DNA loops by cohesin, limited digest with benzonase stripped most cohesin, Pds5b and sororin but not histone H2A off the beads (Figure 1B, lane 7). Chromatin beads washed with high salt were then used as substrate in the prophase pathway reconstitution assay. Confirming our earlier experiments, the removal of Rad21, Pds5b (and sororin) from these chromatin beads required Wapl, Nek2a, Cdk1/2-cyclin A2 and aurora B-INCENP (Figure 1C, lanes 1-5, 7 and 10). In contrast and as seen before, the omission of Plk1 or Cdk1-cyclin B1 did not compromise the release of cohesin and associated factors (Figure 1C, lanes 6 and 8). Heterochromatin protein 1 (HP1) was associated with our beads and reported to recruit Sgo1 to interphase centromeres (Kang, Chaudhary et al., 2011). However, chemical inhibition of PP2A did not increase the efficiency of cohesin liberation (Figure 1C, lanes 9+10) suggesting that the potential recruitment of PP2A by HP1-Sgo1 has no effect on our assay. Interestingly, the kinases were able to selectively solubilize some sororin in the absence of Wapl (Figure 1C, lane 3). This is consistent with a phosphorylation-dependent weakening of the Pds5-sororin interaction (Dreier et al., 2011, Liu et al., 2013a, Nishiyama et al., 2013). Collectively, our reconstitution experiments confirm aurora B as a prophase pathway component. Furthermore, with Nek2a and Cdk1/2-cyclin A2, they identify two kinases that were not previously known to be required for the proteolysis-independent release of cohesin from early mitotic chromatin.

**Figure 1.**
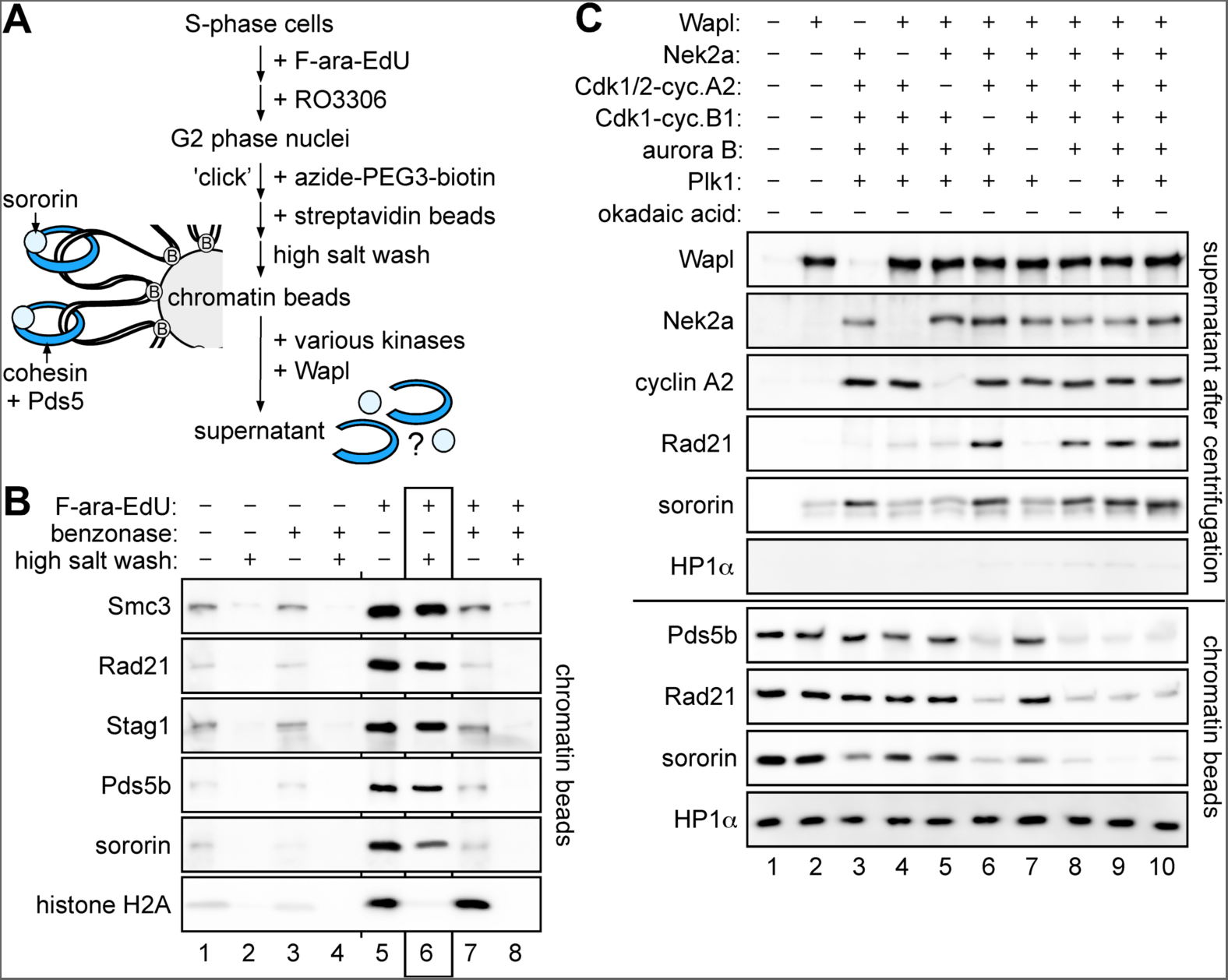
*In vitro* reconstitution of the prophase pathway. **A** Experimental outline. DNA-mediated chromatin pull-down from G2-arrested HeLaK cells is followed by cohesin eviction experiments. **B** Cohesin is eluted from immobilized chromatin upon DNA-cleavage but not high salt treatment. Chromatin beads were treated with benzonase and/or high salt and retained proteins were detected by immunoblotting. High-salt washed chromatin (lane 6) was used for subsequent cohesin eviction experiments. **C** Wapl and the three kinases Nek2a, Cdk1/2-cyclin A2 and Aurora B-INCENP are required for displacement of cohesin from G2-phase chromatin. Immobilized chromatin was combined with the indicated proteins. Following centrifugation, DNA-beads and supernatant were analyzed by immunoblotting.

### Nek2a and Cdk1/2-cyclin A2 phosphorylate Pds5b in its unstructured C-terminal part

To reveal putative targets of Nek2a and Cdk1/2-cyclin A2, we conducted kinase assays. Purified Nek2a phosphorylated *in vitro* expressed Rad21, Smc3, and Pds5b but not Smc1α or Stag1/2, whereas Cdk1/2-cyclin A2 labeled only Pds5b (Figure EV2). We focussed on Pds5b because it is specifically required for centromeric cohesion and modified by Cdk1/2-cyclin A2 on Ser1161 and -1166 *in vivo* (Carretero, Ruiz-Torres et al., 2013, Dumitru, Rusin et al., 2017). In contrast to wild type Pds5b (WT), Pds5b-Ser1161,1166Ala (2A) was resistant to phosphorylation by Cdk1/2-cyclin A2 (Figure EV2, lane 34), arguing that these two residues represent the primary target sites for this kinase within Pds5b. Inspection of nearby sequence stretches for putative Nek2a phosphorylation sites (L/M/F-x-x-S) suggested that Ser1177,-1182, and -1209 of Pds5b might be substrates (van de Kooij, Creixell et al., 2019). Indeed, recombinant Nek2a modified *in vitro*-expressed Pds5b-WT and Pds5b-2A but not Pds5b-Ser1177,1182,1209Ala (3A) (Figure 2A, lanes 1+2, 11+12 and 15+16), while Cdk1/2- cyclin A2 modified Pds5B-WT and -3A but not -2A (Figure 2A, lanes 3+4, 13+14 and 17+18). Consistently, Pds5b-Ser1161,1166,1177,1182,1209Ala (5A) was resistant against phosphorylation by both kinases (lanes 19-22). *In vivo*-phosphorylation of Pds5b on Ser1177 and -1182 was previously reported (Daub, Olsen et al., 2008, Olsen, Vermeulen et al., 2010). To investigate phosphorylation of Ser1209, we raised a phosphorylation specific antibody. It detected Nek2a-phosphorylated Pds5b-WT but not Cdk1/2-cyclin A2-phosphorylated Pds5b- WT or Nek2a-treated Pds5b-3A (Figure EV3A), demonstrating that S1209 is targeted by Nek2a *in vitro*. Anti-pS1209 immunoprecipitated endogenous Pds5b from early mitotic-cells but neither from G2- or prometaphase-cells nor from early mitotic-cells treated with a Nek2a-specific inhibitor (Figure EV3B, lanes 1-4 and 11-25) (Lebraud, Coxon et al., 2014). Thus, *in vivo*-phosphorylation of Ser1209 commences only in early mitosis and in a Nek2a- dependent manner. Moreover, S1209-phosphorylated Pds5b was in the soluble, cytoplasmic fraction and associated with chromatin only in Wapl-depleted prophase cells (Figure EV3B, compare lanes 16+17 with 31+32). This is consistent with Ser1209-phosphorylation of Pds5b promoting the Wapl-dependent release of cohesin from chromosomes.

**Figure 2.**
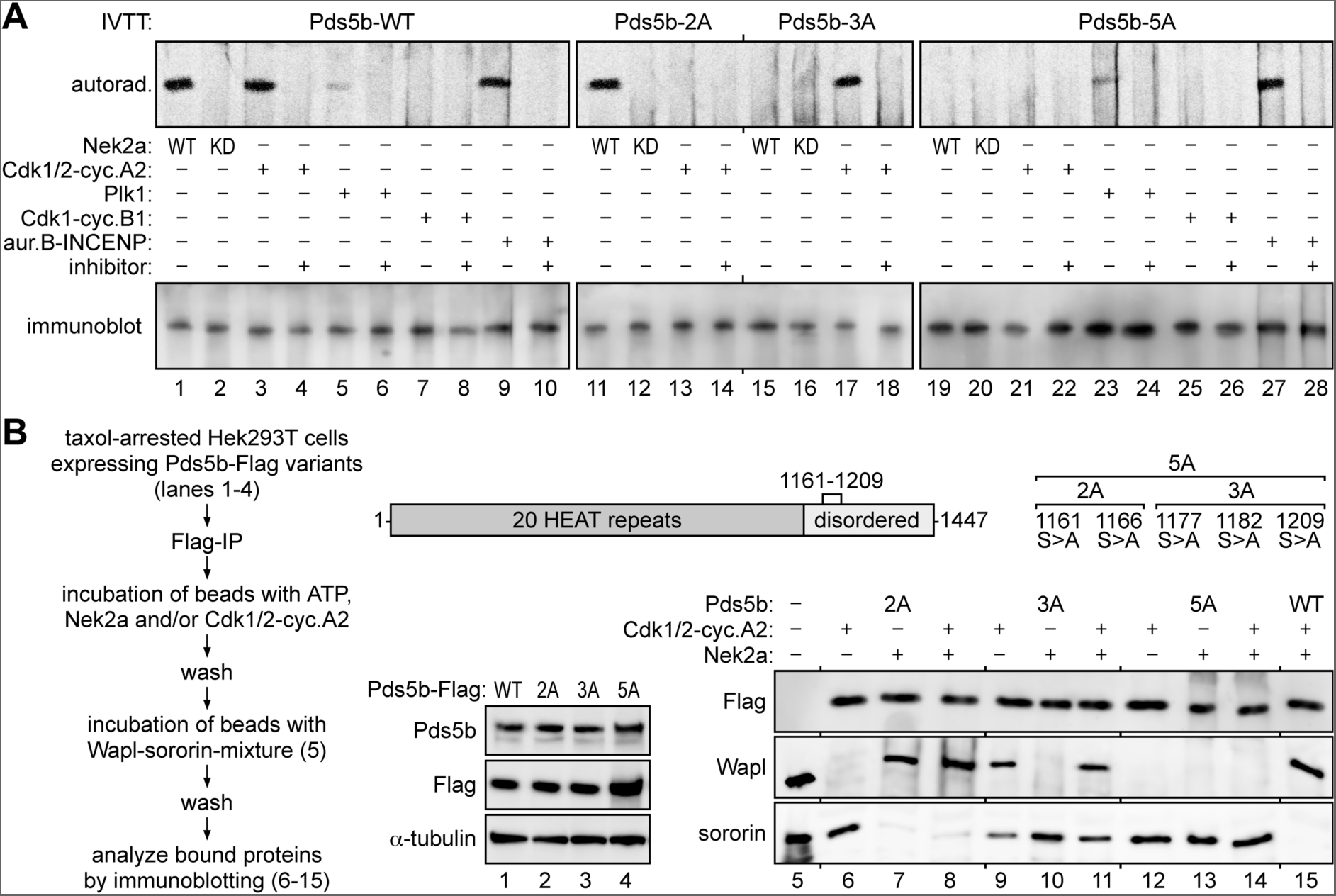
Phosphorylations by Nek2a and Cdk1/2-cyclin A2 turn Pds5b from a sororin- into a Wapl-binder. **A** Nek2a and Cdk1/2-cyclin A2 target juxtaposed but distinct residues within the disordered, C-terminal domain of Pds5b. *In vitro* expressed Pds5b variants were incubated with the indicated kinases and inhibitors in presence of [γ-^33^P]-ATP, subjected to SDS-PAGE and analyzed by autoradiography and immunoblotting. Note that Pds5b is also phosphorylated by aurora B and, weakly, by Plk1. KD = kinase-dead Nek2a-Lys37Met; IVTT = coupled *in vitro* transcription-translation. **B** Rendering Pds5b resistant against both Nek2a and Cdk1/2-cyclin A2 is necessary and sufficient to switch it from a Wapl- to a sororin-binder. Shown are the experimental outline and corresponding immunoblots. Note that following the incubation with Nek2a and/or Cdk1/2-cyclin A2, ATP was no longer present to exclude phosphorylation of sororin (and/or Wapl) by traces of kinases, which might be retained on the beads.

### Phosphorylation of its C-terminal part convert Pds5b from a sororin to a Wapl binder

To study the effect of phosphorylation on Pds5 interactions, Flag-tagged Pds5a or -b were expressed in Hek293T cells either alone or together with stabilized truncation variants of Nek2a (Nek2aΔMR) and cyclin A2 (Δ86-cyclin A2). Following a taxol arrest, Pds5 was affinity-purified from chromatin-free cell lysates. The Pds5-loaded Flag beads, which were free of endogenous Wapl and sororin, were then incubated with a mixture of recombinant Wapl and sororin before bound and unbound fractions were analyzed by immunoblotting (Figure EV4). Importantly, expression of Nek2aΔMR and Δ86-cyclin A2 each improved the binding of Wapl to Pds5b at the expense of sororin. Co-expression of both switched Pds5b almost fully from a sororin- to a Wapl-specific interactor (Figure EV4, lanes 1-4) but had no effect on the association of Pds5a with Wapl and sororin (Figure EV4, lanes 5 and 6). In a variation of this experiment, Flag-tagged Pds5b variants were overexpressed (Figure 2B, lanes 1-4), affinity-purified after an extended prometaphase arrest from Nek2a- and cyclin A2-free cells and only then incubated with recombinant Nek2a and/or Cdk1/2-cyclin A2 in presence of ATP. As before, the Flag-beads were then washed, incubated with recombinant Wapl and sororin (Figure 2B, lane 5), washed again and finally analyzed by immunoblotting for associated proteins (Figure 2B, lanes 6-15). Confirming previous kinase and competition assays, Pds5b- 2A only bound sororin, even after exposure to Cdk1/2-cyclin A2, but preferred Wapl upon phosphorylation by Nek2a (Figure 2B, lanes 6-8). *Vice versa*, the sororin bias of Pds5b-3A was shifted towards Wapl by Cdk1/2-cyclin A2 but not Nek2a (Figure 2B, lanes 9-11). Importantly, Pds5b-5A constitutively bound only sororin, even after exposure to Nek2a and Cdk1/2-cyclin A2 (Figure 2B, lanes 12-14), while Pds5b-WT treated with these two kinases was an exclusive Wapl-interactor (Figure 2B, lane 15). Taken together, these data demonstrate that phosphorylation of Ser1161 and -1166 by Cdk1/2-cyclin A2 and of Ser1177, -1182 and -1209 by Nek2a converts Pds5b from a sororin- to a Wapl-binder. In the context of chromatin, this means Pds5b-dependent recruitment of Wapl to cohesin, thereby providing a mechanistic explanation of how these two kinases promote dissolution of arm cohesion.

### Non-degradable cyclin A2 triggers premature sister chromatid separation

A corollary of Pds5b being a target of prophase pathway signalling is that Sgo1-PP2A must be able to dephosphorylate Pds5b in order to protect pericentromeric cohesion (see below). Consequently, expression of stabilized variants of Nek2a and/or cyclin A2 might prolong the activity of the prophase pathway beyond the time of Sgo1 relocalization and, hence, trigger PCS. Although expression of Nek2aΔMR and Δ86-cyclin A2 trigger apoptosis, this is dependent on separase activation in late metaphase (Hellmuth & Stemmann, 2020). Therefore, HeLaK cells transfected to express Nek2aΔMR and/or Δ86-cyclin A2 were taxol-arrested in prometaphase within the first cell cycle after transfection and then subjected to chromosome spreading (Figure 3A). Indeed, 56(+/-3)% of Δ86-cyclin A2 expressing cells suffered from PCS, while only about 3% of mock-transfected cells exhibited this defect (Figure 3B, columns 1 and 2). Surprisingly, expression of Nek2aΔMR had a much milder effect, causing merely 12% PCS (Figure 3B, column 6; see discussion). Next, we conducted rescue experiments co-expressing Pds5b variants along with the stabilized kinases. Corresponding cells were analyzed by chromosome spreads (Figure 3B) and immunoblotting of cellular contents after their separation into soluble and chromatin fractions (Figure 3C). The Δ86-cyclin A2 triggered PCS phenotype was largely unaffected by Pds5b-WT but almost fully suppressed by Cdk1/2-cyclin A2 resistant Pds5b-2A (Figure 3B/C, columns/lanes 3 and 4). Additionally, this variant strongly inhibited chromosome arm separation and persisted - along with sororin - on isolated mitotic chromatin (column/lane 4). Pds5b-2A (and sororin) even persisted on zipped chromosomes in Nek2aΔMR-expressing cells (Figure 3B/C, column/lane 8), argueing that S1161,1166-phosphorylation might be essential for Wapl-dependent cohesin removal *in vivo*. Conversely, Nek2a-resistant Pds5b-3A (and sororin) did not appreciably accumulate on prophase chromatin of Δ86-cyclin A2 expressing cells (Figure 3C, lane 5). This was surprising because chemical inhibition of Nek2a prevented the release of Rad21, Pds5 and sororin from early mitotic chromatin (Figure EV3B, lanes 2+3, 14+15 and 18+19) and indicated that constitutive S1161,1166-phosphorylation might be sufficient for ring opening. Nevertheless, S1177,1182,1209-phosphorylation of Pds5b clearly contributes to prophase pathway signalling as exemplified by failure of Nek2aΔMR expressing cells to displace Pds5b-3A (and sororin) from chromatin arms (Figure 3B/C, column/lane 9).

**Figure 3.**
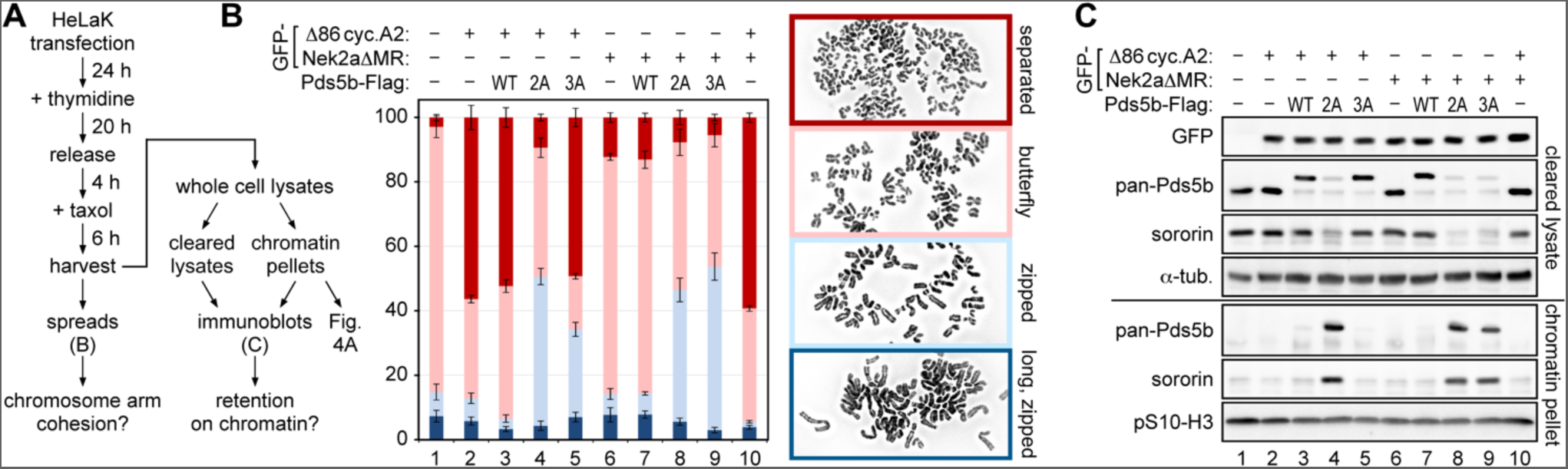
Stabilized Nek2a and cyclin A2 trigger PCS to different extent. **A** Experimental outline. **B** Stabilized cyclin A2 causes profound PCS in prometaphase-arrested cells, which is largely suppressed by Pds5b-2A. Spreads were categorized as separated (= single chromatids), butterfly (= arms separated), zipped (= fully condensed but arms not separated) and long, zipped (early mitotic, not yet fully condensed). Note that Pds5b-2A and -3A partially prevent arm separation as judged by relative increase of zipped chromosomes (light blue). The stacked bars represent the relative percentage (mean value with standard deviation) of each category. For every condition > 100 spreads were counted for each biological replicate (n=3). **C** Sororin is retained by Nek2a- or Cdk1/2-cyclin A2-resistant Pds5b on prometaphase chromatin. Note that i) eGFP-Nek2aΔMR and eGFP-Δ86-cyclin A2 migrate at the same height in SDS-PAGE, ii) expression of transgenic Pds5b results in a lower level of endogenous Pds5b and iii) the retention of Pds5b and sororin on chromatin correlates with the appearance of zipped chromosomes (see B).

### Pericentromeric Pds5b is kept in a de-phosphorylated state

Is pericentromeric Pds5b indeed kept in a de-phosphorylated state? To address this issue, chromatin from prometaphase cells, which expressed Nek2aΔMR and Flag-tagged Pds5b- WT and retained cohesion primarily at pericentromeres of butterfly-shaped, condensed chromosomes (Figure 3B/C, column/lane 7), was used as starting material for a Rad21-IP. Subsequent immunoblotting demonstrated that co-purifying endogenous and Flag-tagged Pds5b-WT were unphosphorylated at S1209 (Figure 4A, lanes 1 and 2). However, S1209- phosphorylation became readily detectable when the Rad21-beads were treated with recombinant Nek2a (in presence of ATP and a PP1/PP2A inhibitor) prior to Western analysis (Figure 4A, lanes 3 and 4). Thus, in prometaphase residual chromosomal Pds5b is unphosphorylated on S1209 - even in presence of stabilized Nek2a. These biochemical experiments were complemented by IF analyses. (Peri)centromeres of isolated prometaphase chromosomes were stained by a pan-specific Pds5b antibody but not by anti- pS1209 (Figure 4B, upper panels). This was not due to the inability of anti-pS1209 to detect phosphorylated Pds5b *in situ* because it labeled mitotic chromosomes from Wapl-depleted cells along their entire length except at (peri)centromeres (lower panels). In conclusion, Pds5b is indeed kept dephosphorylated specifically at (peri)centromeres - at least on Nek2a- targeted Ser1209. Although we did not directly address this here, it is conceivable that the relevant phosphatase is PP2A targeted to pericentromeric cohesin by Sgo1.

**Figure 4.**
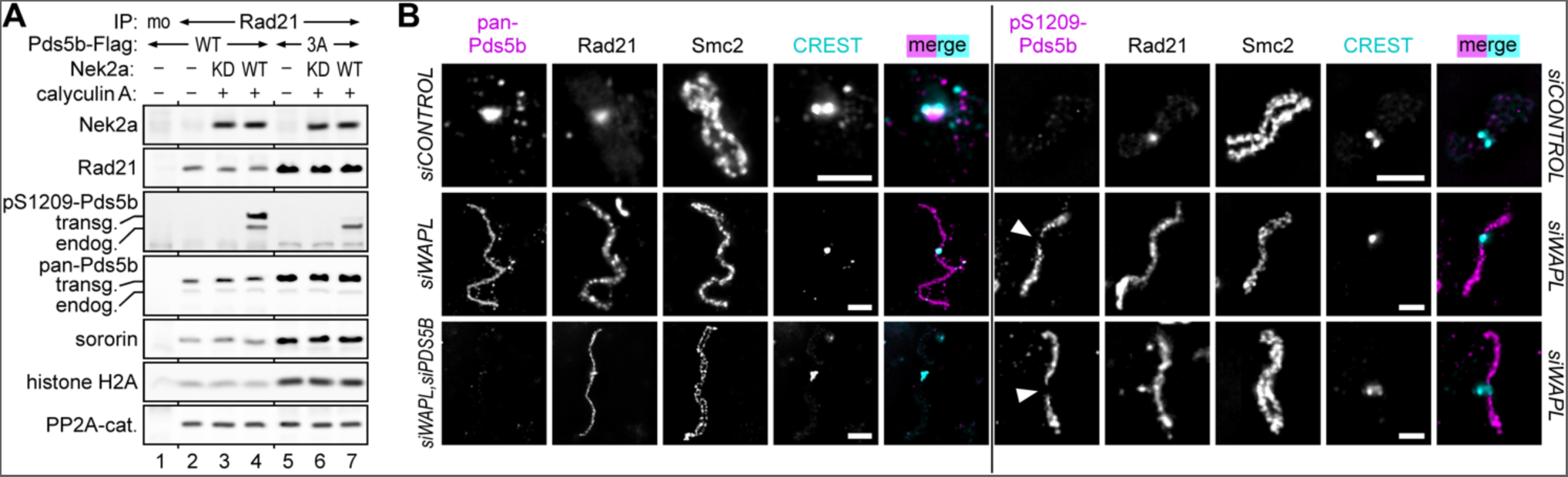
Pericentromeric Pds5b is kept in a dephosphorylated state. **A** Chromatin from Nek2aΔMR- and Flag-tagged Pds5b-WT-expressing cells (see column/lane 7 of figure 3B, C) was benzonase-treated and subjected to Rad21-IP. Precipitated proteins were detected by immunoblotting. Chromatin from Nek2aΔMR- and Flag-tagged Pds5b-3A- expressing cells (see column/lane 9 of figure 3B, C) served as control. mo = mock-IP; KD = kinase-dead; transg. = transgenic; enodg. = endogenous. **B** HeLaK cells were transfected with the indicated siRNAs and released from G2- into a prometaphase arrest. Chromosomes were isolated and subjected to Immunofluorescence microscopy using the indicated antibodies. Arrow heads point at (peri)centromeric regions as judged by CREST staining. Scale bars = 2.5 μm.

### Mimicking Pds5b phosphorylation causes PCS, while preventing it suppresses PCS in absence of Sgo1

What are the cellular consequences of prevented or constitutive Pds5b phosphorylation under more physiological conditions, i.e. without expression of APC/C-resistant Nek2a or cyclin A2? To address this issue, HeLaK cells were transfected to deplete Pds5a/b by RNAi and simultaneously express siRNA-resistant, Flag-tagged Pds5b-variants. In an attempt to mimic constitutive phosphorylation, a new Pds5b-6D (Ser1161,1162,1166,1177,1182,1209Asp) was also included. Corresponding cells were synchronized in prometaphase and subjected to Flag-IP and chromosome spreading (Figure 5). Wild type Pds5b-Flag robustly interacted with Wapl but sororin-binding was undetectable (Figure 5A, lane 1). While Pds5b-6D behaved like wild type (Figure 5A, lane 5), the picture was inverted for Pds5b-2A, -3A and -5A. These Nek2a- and/or Cdk1/2-cyclin A2- resistant variants bound little Wapl but associated with sororin instead (Figure 5A, lanes 2- 4). This exchange of Wapl for sororin, which was most pronounced for Pds5b-5A, correlated with increasing chromatin association as judged by co-IP of histone H2A. Chromosome spreading revealed that most control cells (73%) contained cohesed chromosomes with separated arms and that only few (3%) suffered from PCS (Figure 5B, column 1). Replacing Pds5b-WT by Pds5b-2A, -3A or -5A stepwise increased the fraction of cells containing chromosomes with unseparated/zipped arms from 24% (WT) to 74% (5A) (Figure 5B, columns 2-4). In contrast, 51% of cells expressing Pds5b-6D contained single chromatids (Figure 5B, column 5). Depletion of endogenous Sgo1 by RNAi did not visibly change the interaction behavior of the Pds5b variants (Figure 5A, lanes 6-10), which is consistent with Sgo1-PP2A protecting only a small pool of cohesin. However, and consistent with previous reports (McGuinness et al., 2005, Salic, Waters et al., 2004, Tang et al., 2004), absence of Sgo1 caused PCS in 63% of Pds5b-WT expressing cells (Figure 5B, column 10). While Pds5b- 6D further aggrevated this phenotype (PCS in 75% of cells, Figure 5B, column 6), Ala-containing Pds5b variants increasingly suppressed it, with Pds5b-5A having the strongest rescue effect (PCS in 12% of cells, Figure 5B, column 7). Thus, preventing phosphorylation of Pds5b by Nek2a and Cdk1/2-cyclin A2 renders cohesin partially resistant to Wapl-dependent removal from prophase chromatin. Conversely, constitutive phosphorylation of Pds5b by Nek2a and Cdk1/2-cyclin A2 can be successfully mimicked by Asp-residues and renders the cohesin ring hypersensitive to opening in early mitosis. Mechanistically, these effects can be attributed to Pds5b-5A and -6D binding sororin constitutively or not at all, respectively.

**Figure 5.**
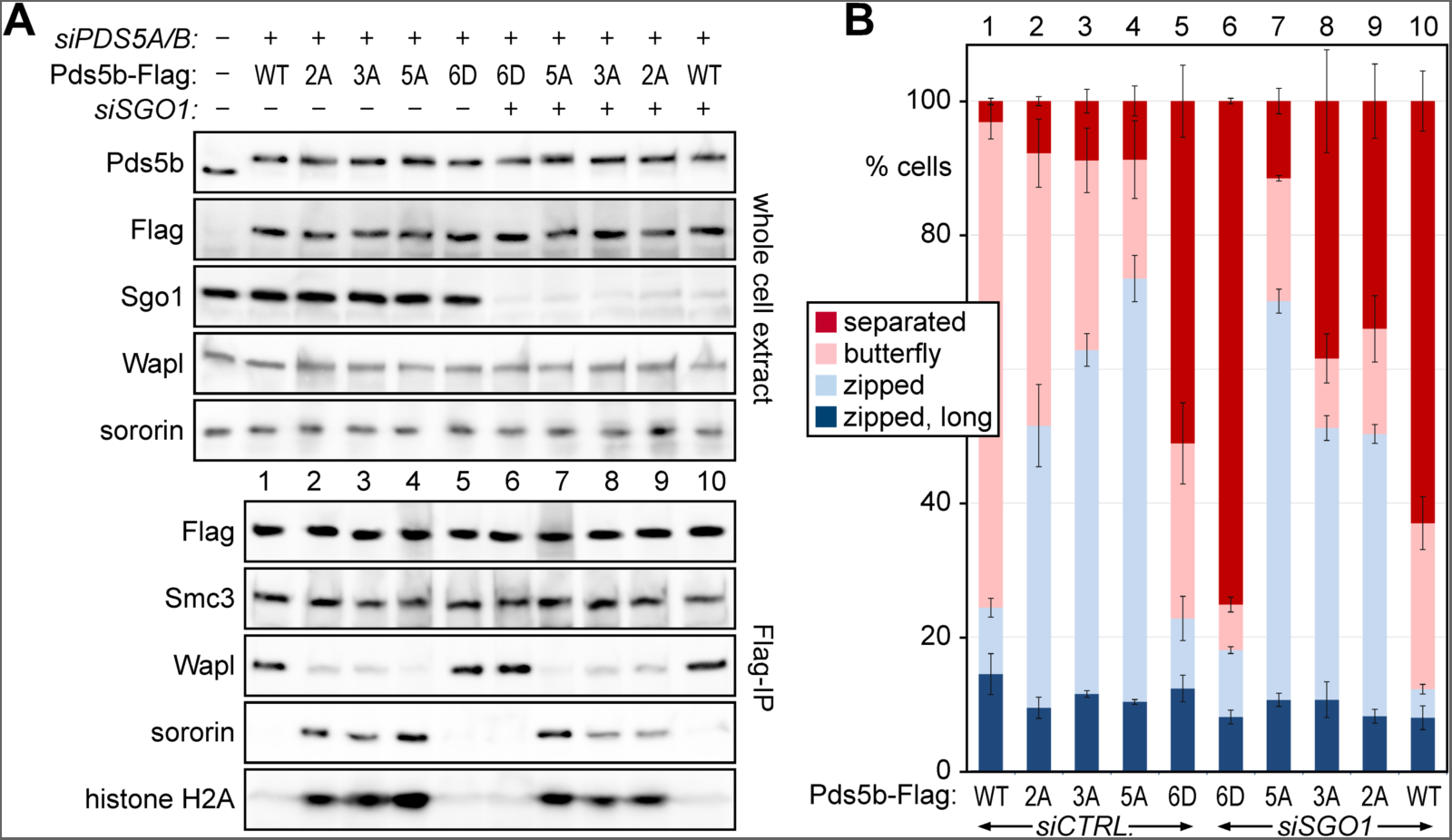
Mimicking Pds5b phosphorylation by Nek2a and Cdk1/2-cyclin A2 causes PCS, while preventing it inhibits arm separation and suppresses PCS in absence of Sgo1. **A** HeLaK cells were transfected to replace endogenous Pds5a/b by Flag-tagged Pds5b variants and, where indicated, to deplete Sgo1. Following synchronization in prometaphase, cells were analyzed by (IP-)Western. **B** Chromosome spreads from cells in A were analyzed as in figure 3B. Note that the co-IP of histone H2A with Pds5b correlates with the abundance of zipped chromosomes, which is consistent with chromosomal retention of cohesin. For every condition > 100 spreads were counted for each biological replicate (n=3).

## Discussion

Here, using purified components, we report the successful reconstitution of the non-proteolytic displacement of cohesin from replicated chromatin. Cohesin release in our system requires only three kinases in addition to Wapl, namely aurora B, Nek2a and cyclin A2-activated Cdk1/2, whereas Plk1 and Cdk1-cyclin B1 are dispensable. The necessity of Nek2a and cyclin A2 strongly suggests that the Wapl-dependent removal of cohesin from chromatin is no longer active by metaphase, when these two factors have already been degraded. This explains why in a normal mitosis there is no Wapl-dependent PCS in the time between deprotection of pericentromeric cohesin (by relocalization of Sgo1-PP2A) until the activation of separase.

The dispensability of Plk1 and Cdk1-cyclin B1 for the *in vitro* release of cohesin from chromatin is surprising because several independent studies have reported the requirement of these two kinases for prophase pathway signaling (Dreier et al., 2011, Gimenez-Abian et al., 2004, Lenart et al., 2007, Liu et al., 2013a, Losada et al., 2002, Nishiyama et al., 2013, Sumara et al., 2002). Cdk1/2-cyclin A2 and Cdk1-cyclin B1 have similar substrate specificities, although in our hands Cdk1/2-cyclin A2 can phosphorylate Pds5b, while Cdk1- cyclin B1 cannot (Figure 2A) (Moore, Kirk et al., 2003). Thus, it is conceivable that Cdk1/2- cyclin A2 can functionally replace Cdk1-cyclin B1 in prophase pathway signaling, but not *vice versa*. The dispensability of Plk1 could be explained if prophase pathway components are activated by this kinase *in vivo* and are added in their already active form to our *in vitro* system. According to the literature, aurora B and Wapl are candidates for putative Plk1- activated prophase pathway components (Challa, Fajish et al., 2019, Rosasco-Nitcher, Lan et al., 2008). We isolate Wapl from mitotically arrested cells, i.e. in a state of high Plk1 activity. Therefore, it will be interesting to test whether its dephosphorylation would be sufficient to establish a Plk1 requirement for cohesin release in our assay.

Cyclin A2-resistant Pds5b-2A (and sororin) accumulated on mitotic chromatin - even in cells expressing Nek2aΔMR (Figure 3C). This suggests that phosphorylation of Pds5b on serines 1161 and 1166 may be essential for Wapl-dependent cohesin removal and cannot be functionally replaced by Nek2a-dependent phosphorylation of nearby sites. Conversely, the accumulation of Nek2a-resistant Pds5b-3A (and sororin) on mitotic chromatin could be prevented by co-expression of Δ86-cyclin A2 (Figure 3C), suggesting that continued phosphorylation of Pds5b on S1161,1166 may be sufficient to compensate for the loss of Nek2a. These interpretations may also explain why Δ86-cyclin A2 expression caused profound PCS, whereas the premature loss of cohesion caused by Nek2aΔMR expression was almost negligible (Figure 3B). Thus, only the stabilization of cyclin A2, but not of Nek2a, seems to prolong the activity of the prophase pathway beyond the time of cohesin deprotection by departure of Sgo1-PP2A. Nevertheless, Nek2a clearly contributes to prophase pathway signaling, as exemplified by profound retention of Rad21, Pds5b and sororin upon chemical inhibition of Nek2a (Figure EV3B) and the ability of Nek2a-resistant Pds5b-3A to partially suppress PCS caused by siRNA-mediated depletion of Sgo1 (Figure 5B).

Nek2a- and Cdk1/2-cyclin A2 phosphorylate Pds5b at 5 sites within a 50 residues window of the long (330 amino acids), unstructured C-terminal part. How exactly these phosphorylations convert Pds5b from a sororin to a Wapl binder, we cannot yet say. However, it is interesting to note that Nek2a, Cdk1/2-cyclin A2 and aurora B are able to selectively elute sororin (without cohesin) from immobilized chromatin in the absence of Wapl (Figure 1C, lane 3). This suggests that the primary effect of phosphorylation is the weakening of sororin’s interaction with Pds5-cohesin rather than the enforcement of Wapl’s interaction with these partners. With their shared YSR- and FGF-motifs, sororin and Wapl compete for the same contact sites on Pds5, but AlphaFold predicts an additional interaction of sororin with the Smc3-Rad21 interface, which is specific, i.e. not shared by Wapl (Nasmyth et al., 2023). As it is difficult to envision how phosphorylation of Pds5b’s C-terminal part could have different effects on very similar interactions, we favor the idea that it may instead interfere with the interaction of sororin with cohesin’s DNA exit gate.

The phosphorylation sites within Pds5b targeted by Nek2a- and Cdk1/2-cyclin A are not conserved in Pds5a. The serines are either absent (positions 1161, 1177, 1209) or in an altered sequence context that no longer matches the substrate consensus sequences of the two kinases (positions 1166 and 1182). Accordingly, the binding of sororin or Wapl to Pds5a is not modulated by Nek2a or Cdk1/2-cyclin A2 (Figure EV4). This may imply that these two kinases are dispensable for the Wapl-dependent release of Pds5a-bound cohesin. Given that pericentromeric cohesion depends on Pds5b (Carretero et al., 2013), this would not contradict our proposal that once Sgo1 has left cohesin, Wapl can no longer displace it. In fact, such a difference in regulation may even be the reason for the existence of two Pds5s in vertebrates. Alternatively, Pds5a-mediated cohesion could also be sensitive to Nek2a- and Cdk1/2-cyclin A2. Cohesin is not eluted from chromatin *in vitro* when exposed to Wapl, Plk1, Cdk1-cyclin B1 and aurora B, but most of Rad21, Smc3 and sororin is solubilized when Nek2a and Cdk1/2-cyclin A2 are additionally included (Figures 1C and EV1). This, together with the profound chromosomal retention of Rad21 and sororin upon chemical inhibition of Nek2a *in vivo* (Figure EV3B), actually favors this second possibility. If Pds5a-cohesin is indeed also responsive to Nek2a- and Cdk1/2-cyclin A2, then there would have to be additional substrates of these kinases in the prophase pathway, with sororin being an obvious candidate.

## Methods

### Cell lines and plasmid transfections

Human cells (HeLaK and Hek293T) were cultured in DMEM (Biowest) supplemented with 10% FCS (Sigma-Aldrich) at 37°C and 5% CO_2_. pCS2-based expression plasmids were transfected into HeLaK cells with Lipofectamine 2000 (Invitrogen) according to manufacturer’s protocol and into Hek293T cells using a calcium phosphate-based method. Pds5a/b variants were expressed in fusion with a C-terminal Tev2-Flag3 tag, while Nek2aΔMR, Δ86-cyclin A2 and Wapl were expressed in fusion with an N-terminal GFP- SUMOstar tag.

### RNA interference

*siRNAs: PDS5B*: 5’-GGAUAGAUCUUAAGCAGUATT-3’,

*PDS5A_1*: 5’- GGGAAAGAACACUGGAUAATT-3’,

*PDS5A_2*: 5’- UGUAAAAGCUCUCAACGAATT-3’,

*WAPL_1*: 5’- CGGACUACCCUUAGCACAA -3’,

*WAPL_2*: 5’- GGUUAAGUGUUCCUCUUAUTT -3’,

*SGO1_1:* 5’- AGUAGAACCUGCUCAGAA -3’,

*SGO1_2:* 5’- GAUGACAGCUCCAGAAAUUTT -3’,

*LUCIFERASE siRNA* served as negative control *(siCONTROL)*. HeLaK and Hek293T cells were transfected with with 50-100 nmol siRNA duplex (Eurofins Genomics) using RNAiMax (Invitrogen) or a calcium phosphate-based method, respectively. To increase target knockdown efficiency siRNAs 1 and 2 were transfected simultaneously. Since depletion of Sgo1 causes premature loss of cohesion, mitotic arrest and cell death, cells were transfected approximately halfway through of an 18-20 h thymidine block, then released and analyzed in the following mitosis. In all other cases, siRNA transfer was performed before synchronization procedures were applied.

### Cell cycle synchronizations

For synchronization in early S-phase, cells were treated with thymidine (2 mM; Sigma-Aldrich) for 20 h. For experiments with ectopic expression of Pds5 variants, plasmid transfection was done 8-12 h prior to thymidine addition. Synchronization of cells in prometaphase was achieved by 90 min-incubation with taxol (0.2 μg/ml; LC Laboratories) 9 h after release from a thymidine block. Alternatively, i.e. without pre-synchronization with thymidine, taxol was added for 10 h to asynchronous cells 36 h after plasmid transfection. Transfected, prometaphase-arrested cells were analyzed by chromosome spreading and served as source of chromatin-free Pds5a/b for Wapl/sororin binding assays. For an arrest in late G2 phase, cells released for 6 h from a thymidine block were supplemented with RO- 3306 (8-10 µM; Santa-Cruz Biotechnology) for a maximum of 6 h. To visualize early mitotic events, RO-3306 was washed out with pre-warmed media, and cells were fixed 20-30 min thereafter. Alternatively, G2 cells were released into taxol for 2 h to re-arrest them in prometaphase.

### Antibodies

Non-commercial antibodies used in this study were generated at Charles River Laboratories. To this end, antigenic peptides (Bachem) were coupled via terminal Cys to Maleimide activated KLH (ThermoFisher) prior to immunization. Antibodies were affinity purified against immobilized antigens. Antigenic proteins were coupled to NHS-activated sepharose (GE Healthcare), whereas antigenic peptides were coupled to SulfoLink coupling gel (ThermoFisher). For immunoblotting the following antibodies diluted in 1xPBS, 1% (w/v) BSA (Roth) were used: Rabbit anti-Smc1 (1:1,000; A300-055A, Bethyl), rabbit anti-Smc3 (1 µg/ml) (Schockel, Mockel et al., 2011), guinea pig anti-Rad21 (1.8 µg/ml; raised against A107-S271 of the human protein), mouse anti-Rad21 (1:800; B-2, Santa Cruz Biotechnology), rabbit anti-Rad21 (1:1,000; A300-080A, Bethyl), rabbit anti-Stag1 (3,5 µg/ml; raised against the C-terminal 199 amino acids of *Xenopus laevis* SA1; kindly provided by Susannah Rankin, University of Oklahoma, USA) and rabbit anti-Stag2 (3.3 µg/ml; kindly provided by Jan-Michael Peters, Institute of Molecular Pathology, Austria), guinea pig anti-*p*S1209-Pds5b (0.25 µg/ml; raised against CDLVR*p*SELEK = Cys + amino acids 1205-1213 of the human protein, reactivity against the unphosphorylated peptide was removed), guinea pig anti-pan- Pds5b (1.5 µg/ml; raised against P1137-M1308 of the human protein), mouse anti-cyclin A2 (1:200; 46B11, Santa Cruz Biotechnology), mouse anti-Nek2 (1:600; clone 20, 610549, BD Transduction Laboratories), mouse anti-sororin (1:1,000; sc-365319, Santa Cruz Biotechnology), rabbit anti-sororin (1.7 µg/ml) (Wolf, Cuba Ramos et al., 2018), guinea pig anti-Wapl (1.2 µg/ml; raised against M1-C625 of the human protein), rabbit anti-Sgo1 (1:500; ab21633, Abcam), guinea pig anti-Sgo1 (2.5 µ/ml; raised against S105-K281 of the human isoform 1), mouse anti-PP2A-C (1:1,000; 1D6, Millipore), mouse anti-HP1 (1:1,000; hybridoma supernatant PCRP-CBX5-2D8 from DSHB, University of Iowa, USA), rabbit anti-histone H2A (1:1,000; 11-7017, Abeomics), rabbit anti-pS10-histone H3 (1:1,000; 06-570, Merck), mouse anti-α-tubulin (1:200; hybridoma supernatant 12G10; from DSHB, University of Iowa, USA), mouse anti-Flag (1:2,000; M2, Sigma-Aldrich) and mouse anti-GFP (1:2,000; hybridoma supernatant; clone 71; gift from D. van Essen and S. Saccani). Secondary antibodies for immunoblotting were horseradish peroxidase (HRP)-conjugated goat anti-rabbit, anti-mouse and anti-guinea-pig IgGs (1:20,000; Sigma Aldrich). For immunoprecipitation experiments, the following affinity matrices and antibodies were used: Mouse anti-Flag M2-Agarose (Sigma-Aldrich), anti-GFP nanobody covalently coupled to NHS-agarose (GE Healthcare) and guinea pig anti-Rad21 and guinea pig anti-Pds5b coupled to protein A sepharose (GE Healthcare). For non-covalent coupling of antibodies to beads, 10 μl of protein A sepharose were rotated with 2-5 μg antibody for 90 min at room temperature and then washed three times with 1xPBS, 1 % BSA (w/v; Roth). For immunofluorescence microscopy, the following antibodies were used: Guinea-pig anti-pan- Pds5b and anti-pS1209-Pds5b (each at 1.5 µg/ml), mouse anti-Rad21 (1:500; 05-908, Millipore), rabbit anti-Smc2 (2 µg/ml; raised against CAKSKAKPPKGAHVEV = Cys + amino acids 1183-1197 of the human protein) and human anti-CREST (1:1,000; hct-0100, ImmunoVision). Secondary antibodies (all 1:500): Cy3 donkey anti-guinea pig IgG and Cy3 goat anti-rabbit IgG (Invitrogen); Alexa Fluor 488 goat anti-rabbit IgG and Cy5 goat anti-human IgG (Bethyl); Alexa Fluor 350 goat anti-mouse and Alexa Fluor 488 goat anti-mouse IgG (Invitrogen).

### Immunoprecipitation (IP)

2×10^7^ cells were lysed with a dounce homogenizer in 2 ml lysis buffer 2 [20 mM Tris-HCl, pH 7.7; 100 mM NaCl; 10 mM NaF; 20 mM β-glycerophosphate; 10 mM MgCl_2_; 0.1% Triton X- 100; 5% glycerol; 1x complete protease inhibitor cocktail (Roche)], and incubated on ice for 10 min. To preserve phosphorylations, lysis buffer 2 was additionally supplemented with calyculin A (50 nM, LC-Laboratories). Corresponding whole cell lysates were centrifuged for 20 min at 4°C and 16,000 x g to fractionate them into cleared lysates (soluble supernatant) and pelleted chromatin. For IP of soluble proteins, 10 μl of antibody-loaded beads were rotated with 2 ml cleared lysate for 4-12 h at 4°C, washed 5x with lysis buffer 2 and equilibrated in corresponding reaction buffers. For IP of DNA-associated proteins, pelleted chromatin was washed twice in lysis buffer 2 and digested for 1 h at 4°C with benzonase (30 U/l; Santa-Cruz Biotechnology) in 2 ml lysis buffer 2. Then, insoluble material was removed by centrifugation (5 min at 4°C and 2,500 x g), and the supernatant was combined with 10 μl of antibody-loaded beads and rotated for 4-12 h at 4°C. Thereafter, beads were washed 6x with lysis buffer 2 supplemented with additional 250 mM NaCl before they were used for further applications.

### Immunofluorescence microscopy of spread chromosomes

HelaK transfected with the indicated siRNAs were synchronized in G2-phase by addition of RO-3306 followed by subsequent release into taxol. Prometaphase-arrested cells were collected by shake-off 30 min thereafter and immediately processed. Following their pelleting (300 x g, for 3 min), cells were resuspended in hypotonic buffer I (50 mM sucrose, 30 mM TRIS-HCl ph 8.2, 17 mM sodium citrate trihydrate, 0.2 µg/ml taxol), incubated for 7 min, sedimented again and resuspended in hypotonic buffer II (100 mM sucrose). Immediately thereafter, 5 µl cell suspension were dropped onto a coverslip wetted by immersion in fixation buffer (1% PFA, 5 mM sodium borate pH 9.2, 0.15 % TritonX-100) and spread by tilting of the coverslip. After drying at RT, coverslips were washed 3x with PBS, blocked for 1 h at RT in PBS; 3% (w/v) BSA and incubated with primary antibodies for 1 h in a wet chamber. Coverslips were then washed 4x with PBS-Tx (PBS; 0.1% Triton X-100), incubated with fluorescently labelled secondary antibodies for 40 min, washed again 4x and finally mounted in 20 mM Tris-HCl, pH 8.0; 2,33% (w/v) 1,4-diazabicyclo(2.2.2)octane; 78% glycerol on a glass slide. Samples were analyzed on a DMI 6000 inverted microscope (Leica) using a HCX PL APO 100x/1.40-0.70 oil objective collecting Z-stacks series over 2 µm in 0.2 µm increments. Images were deconvoluted and projected into one plane using the LAS-AF software. Smc2 staining was used to identify mitotic chromosomes. Standard chromosome spreads were done as described (McGuinness et al., 2005) and observed on a Axioplan 2 microscope (Zeiss) using a Plan-APOCHROMAT 100x/1.40 oil objective. A cell was counted as suffering from PCS when > 50% of its chromosomes were separated into single chromatids and as defective in prophase pathway signaling when > 50% of its chromosomes exhibited zipped arms. At least 200 spreads were counted per condition.

### Isolation of crude chromatin

Crude G2 chromatin was freshly prepared for each experiment according to a protocol by Mendez and Stillman (Mendez & Stillman, 2000). Briefly, 4 x 10^7^ G2-arrested HelaK cells were resuspended in 3 ml buffer A [10 mM HEPES, pH 7.9; 10 mM KCl; 1.5 mM MgCl_2_; 0.34 M sucrose; 10 % glycerol; 1 mM DTT; 1x EDTA-free complete protease inhibitor cocktail (Roche)] and lysis was initiated by addition of TritonX-100 to 0.1 %. Following a 5 min incubation on ice, chromatin was pelleted by centrifugation for 4 min at 4°C and 1,300 x g, washed once in TritonX-100-supplemented (0.1%) buffer A, re-isolated and resuspended in 3 ml buffer B (3 mM EDTA; 0.2 mM EGTA; 1 mM DTT; 1 mM spermidine; 0.3 mM spermine; 1x complete protease inhibitor cocktail). After incubation for 25 min on ice, the chromatin was sedimented for 4 min at 4°C and 1,700 x g, washed once in 1 ml buffer B and finally resuspended in 200 µl chromatin buffer (5 mM Pipes, pH 7.2; 5 mM NaCl; 5 mM MgCl_2_; 1 mM EGTA; 10 % glycerol).

### DNA-mediated chromatin pull-down

The following procedure is a combination of published protocols (Aranda et al., 2019, Kliszczak et al., 2011, Neef & Luedtke, 2011). Approximately 20 x 10^7^ HeLaK cells pre-synchronized by thymidine treatment (2 mM for 18 h) were released to undergo S-phase in presence of F-ara-Edu (5 µM; Sigma-Aldrich) for 5 h. Then, RO-3306 (10 µM) was added and the G2-arrested cells were harvested and PBS-washed 5 h later. The cell pellet was immediately resuspended in 20 volumes lysis buffer 1 [20 mM HEPES, pH 7.5; 0.22 M sucrose; 2 mM MgCl_2_; 0.5 % (v/v) Nonidet P-40; 0.5 mM DTT; 1x EDTA-free complete protease inhibitor cocktail (Roche)], dounce homogenized and incubated for 10 min on ice. The whole cell lysate was then centrifuged for 15 min at 4°C and 3000 x g in a swing-out rotor to pellet G2 chromatin/nuclei. After removal of the supernatant, the chromatin pellet was supplemented (in this order) with (+)-sodium-L-ascorbate (10 mM; Sigma-Aldrich), azide-PEG3-biotin (0.1 mM; Sigma-Aldrich) and cooper(II)sulfate (2 mM; Roth) and slowly rotated in the dark for 30 min at room temperature (RT). This was followed by three consecutive wash steps (15 min at 4°C and 3,000 x g in a swing-out rotor) with 1 ml lysis buffer 1 each to remove unbound biotin. To increase purity, resuspended chromatin was centrifuged through a 3 cm cushion (20 mM HEPES, pH 7.5; 2 mM MgCl_2_; 0.34 M sucrose) for 20 min at 4°C and 3,400 x g in a swing-out rotor. The resulting chromatin pellet was finally resuspended in 300 µl lysis buffer. High capacity streptavidin beads (250 µl; Thermo Scientific) were pre-blocked with 0.4 mg/ml sonicated salmon sperm DNA in lysis buffer 1 for 30 min at RT, washed twice (centrifugation: 1 min at 4°C and 120 x g) and then combined with the biotinylated G2 chromatin in a final volume of 4 ml. Where indicated, benzonase (30 U/l; Santa Cruz Biotechnology) was added. After 40 min of rotating at 4°C, beads were washed once with lysis buffer 1 and then resuspended in cold high salt buffer (HSB; 20 mM HEPES, pH 7.5; 0.6 M NaCl; 2 mM MgCl_2_; 0.22 M sucrose; 0.2 % NP-40; 1 mM DTT). Following a 10 min incubation, the salt-extracted chromatin was very carefully washed twice with HSB and twice with lysis buffer (using cut-off tips). Finally, the chromatin-loaded streptavidin beads were resuspended in 1 ml nuclear storage buffer [20 mM TRIS-HCl, pH 8.0; 75 mM NaCl; 0.5 mM EDTA; 50 % (v/v) glycerol; 1mM DTT; 1x EDTA-free complete protease cocktail (Roche)] and stored at -20°C.

### Recombinant protein expression and purification

Active Cdk1-cyclin B1 was expressed in insect cells and purified as described (Gorr et al, 2005). Active Plk1 was purchased from R&D Systems (No. 3804-KS). To generate active aurora B kinase, the complex of co-expressed fragments of human aurora B kinase (amino acids 65-344) and INCENP (amino acids 819-918) was affinity purified from bacteria essentially as described (Sessa, Mapelli et al., 2005). Active Nek2a (active or kinase-dead, KD = Lys37Met) and Cdk1/2-cyclin A2 were expressed and purified as described (Hellmuth & Stemmann, 2020). Prior to use, all kinases were tested for activity using 2 µg each of histone H1 (NEB) or Myelin Basic Protein (MBP, Upstate Biotechnology) as model substrates. Human Wapl was purified from 5 x 10^7^ transiently transfected Hek293T cells. To enrich the pool of soluble Wapl, cells were treated with taxol 36 h after plasmid transfection and for 12 h to arrest them in prometaphase prior to harvesting. Following lysis of the cells in 10 ml lysis buffer 2 and clearing of the lysate by centrifugation (30 min at 4°C and 18,000 x g), GFP- SUMOstar-Wapl was immunoprecipitated for 90 min with 50 µl anti-GFP nanobody beads. Then, the beads were washed 5x with lysis buffer 2, transferred into cleavage buffer [10 mM Hepes-KOH, pH 7.7; 50 mM NaCl; 25 mM NaF; 1 mM EGTA; 20% glycerol] and finally rotated for 30 min at 18°C with 5 µg His_6_-SUMOstar protease. The corresponding eluate (50 µl) was snap frozen in aliquots and stored at -80°C. His_6_-SUMO3-sororin was expressed from a pET28-derivative in *E. coli.* Rosetta 2 DE3 (Novagen). Bacteria were lysed in phosphate buffered high saline (PBHS; 10 mM Na_2_HPO_4_; 2 mM KH_2_PO_4_; 500 mM NaCl; 2.7 mM KCl; 5 mM imidazole; 0.5 mM DTT), purified over Ni^2+^-NTA-Agarose (Qiagen) according to standard procedure and eluted in PBHS supplemented with imidazole to 250 mM and pH-adjusted with HCl to 7.5. Following its dialysis into sororin storage buffer (20 mM Tris-HCl, pH 6.9; 100 mM NaCl), His_6_-SUMO3-sororin was stored in aliquots at -80°C.

### *In vitro* cohesin release assay

To assemble kinase reactions, 15 µl chromatin (crude or immobilized on streptavidin sepharose) equilibrated in kinase buffer [10 mM Hepes-KOH, pH 7.7; 50 mM NaCl; 25 mM NaF; 1 mM EGTA; 20% glycerol; 10 mM MgCl_2_; 5mM MnCl_2_; 5 mM DTT] were combined with ATP (1 mM), okadaic acid (1 µM, Alexis Biochemicals) or calyculin A (20 nM, LC Laboratories) and with 0.4 µg Cdk1-cyclin B1, 0.1 µg aurora B-INCENP, 0.1 µg Plk1, 1 µl Nek2a and/or 1 µl Cdk1/2-cyclin A2, as indicated. The samples, which had a total volume of 25-30 µl, were incubated for 20-30 min at 37°C while slowly rotating. This was followed by addition of RO- 3306 (2 µM, Santa-Cruz), BI-2536 (100 nM, Boehringer-Ingelheim), staurosporine (300 nM, Abcam) and/or ZM-447439 (0.5 µM, Tocris), as indicated. After 10 min at RT, reactions were supplemented with 2 µl Wapl and incubated for 20 min at 37°C while slowly rotating. Then, the chromatin was pelleted for 1 min at 4°C and 1,700 x g and the supernatant was recovered for immunoblotting. The chromatin was washed twice with 1 ml each of lysis buffer 2, re-pelleted for 4 min at 1,700 x g and finally resuspended in 20 µl 2x SDS-sample buffer for analysis.

### Kinase and binding assays

For phosphorylation of FLAG- and Rad21-IPs, 15 µl beads (instead of chromatin) were used to assemble kinase reactions essentially as described above. Samples were incubated for 20- 30 min at 37°C while slowly rotating. If the kinase reactions were directly analyzed by immunoblotting, then the beads were pelleted for 1 min at 4°C and 1,700 x g, washed once with 1 ml lysis buffer 2, re-pelleted and eluted by boiling in 15 µl 2x SDS sample buffer. To assess sororin/Wapl binding to Pds5, FLAG-IPs (directly from cells or after *in vitro*- phosphorylation) were washed twice in cleavage buffer containing kinase inhibitors, resuspended in 15 µl cleavage buffer containing 2 µl Wapl and 0.3 µg sororin and rotated for 20 min at 37°C. Then, the supernatant was recovered for later immunoblotting and the beads were washed twice with 1 ml each of lysis buffer 2 before they, too, were eluted by boiling in SDS-sample buffer. For radioactive kinase assays, Rad21, Smc1α, Smc3, Stag1, Stag2 and Pds5b variants were first expressed by ‘cold’ coupled *in vitro* transcription-translation (IVTT; TNT reticulocyte lysate, Promega). IVTT aliquots (3 µl) were then incubated for 20-30 min at 37°C with γ-^33^P-ATP (40 µCi, Hartmann Analytic), ‘cold’ ATP (5 µM) and the indicated kinases in a total volume of 25-30 µl (see above). To specifically inhibit kinase activity during the incubation at 37°C, the corresponding inhibitor (see above) was added together with the kinase during reaction assembly at 4°C. Following SDS-PAGE, radioactive reactions were blotted onto PVDF-membrane (SERVA) and first analyzed by autoradiography. The same membrane was then re-activated with methanol and further examined by immunoblotting.

## Acknowledgements

We thank B. Neumann for recombinant sororin and aurora B kinase and S. Heidmann, T. Klecker and J. Michl for critical reading of the manuscript. This work was supported by a grant (STE997/8-1) from the Deutsche Forschungsgemeinschaft (DFG) to O.S.

## Author contributions

S.H. carried out all experiments. S.H. and O.S. co-designed the research and wrote the paper.

## Data availability

The authors declare that all data supporting the findings of this study are available within the paper and its supplementary information files. If there is reasonable request data can also be provided from the corresponding author.

## Expanded View Figures and legends

**Figure EV1.**
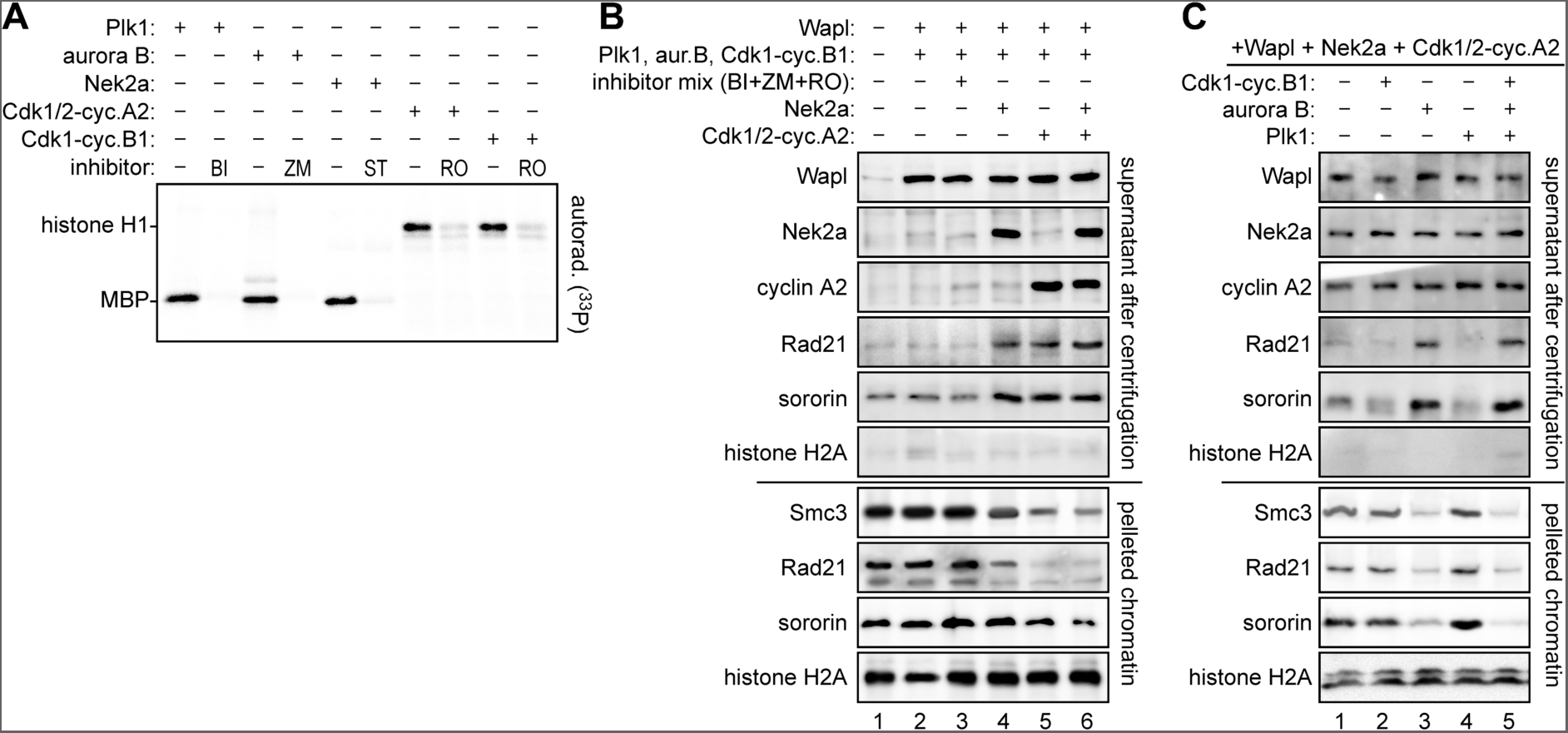
Reconstitution of Wapl-dependent displacement of cohesin from crude G2- phase chromatin. **A** Recombinant kinases are active as judged by phosphorylation of model substrates. Plk1, aurora B-INCENP, Nek2a, Cdk1/2-cyclin A2 or Cdk1-cyclin B1 supplemented with their specific inhibitor or carrier solvent DMSO (-) were incubated with corresponding model substrate in presence of [γ^33^P]-ATP. Reactions were subjected to SDS-PAGE followed by autoradiography. BI, BI2536; ZM, ZM447439; ST, staurosporine; RO, RO3306; MBP, myelin basic protein. **B** Nek2a and Cdk1/2-cyclin A2 are necessary for Wapl-dependent *in vitro* displacement of cohesin from chromatin. Chromatin was freshly purified by repeated sedimentation from G2-arrested HeLaK cells, incubated with recombinant Wapl, kinases and inhibitors (BI2536, ZM447439 and RO3306), as indicated, and then re-pelleted. Supernatants and chromatin pellets were analyzed by immunoblotting. **C** Wapl and the three kinases Nek2a, Cdk1/2-cyclin A2 and Aurora B-INCENP are sufficient to support displacement of cohesin from isolated chromatin. Experiment was done as in B).

**Figure EV2.**
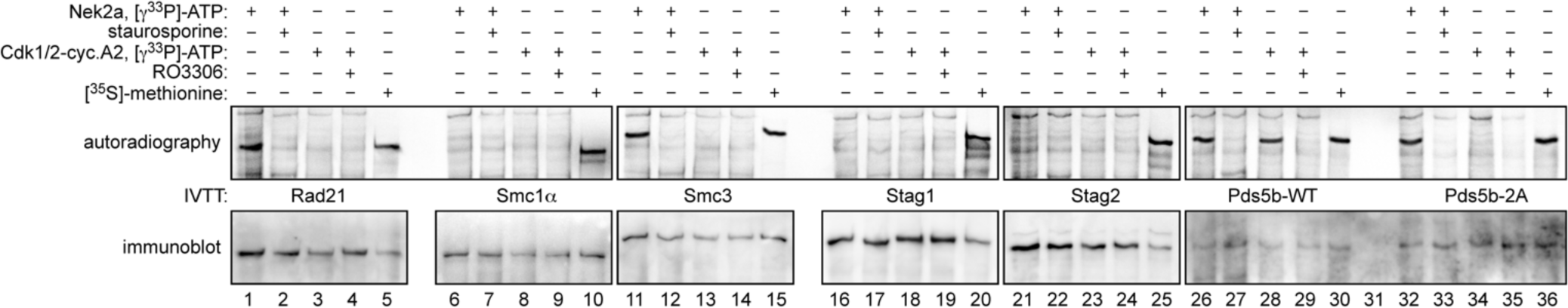
Nek2a and Cdk1/2-cyclin A2 both phosphorylate *in vitro* expressed Pds5b. Cohesin subunits and Pds5b variants were expressed by coupled *in vitro* transcription-translation (IVTT), incubated with Nek2a, Cdk1/2-cyclin A2, [γ-^33^P]-ATP and kinase inhibitors, as indicated, separated by SDS-PAGE and analyzed by autoradiography and immunoblotting. ^35^S-methionine labeled IVTT products served as indicators of the migration behavior of the respective full-length protein. Note that Pds5b-2A (Ser1161,1166Ala) is resistant to phosphorylation by Cdk1/2-cyclin A2 but not Nek2a.

**Figure EV3.**
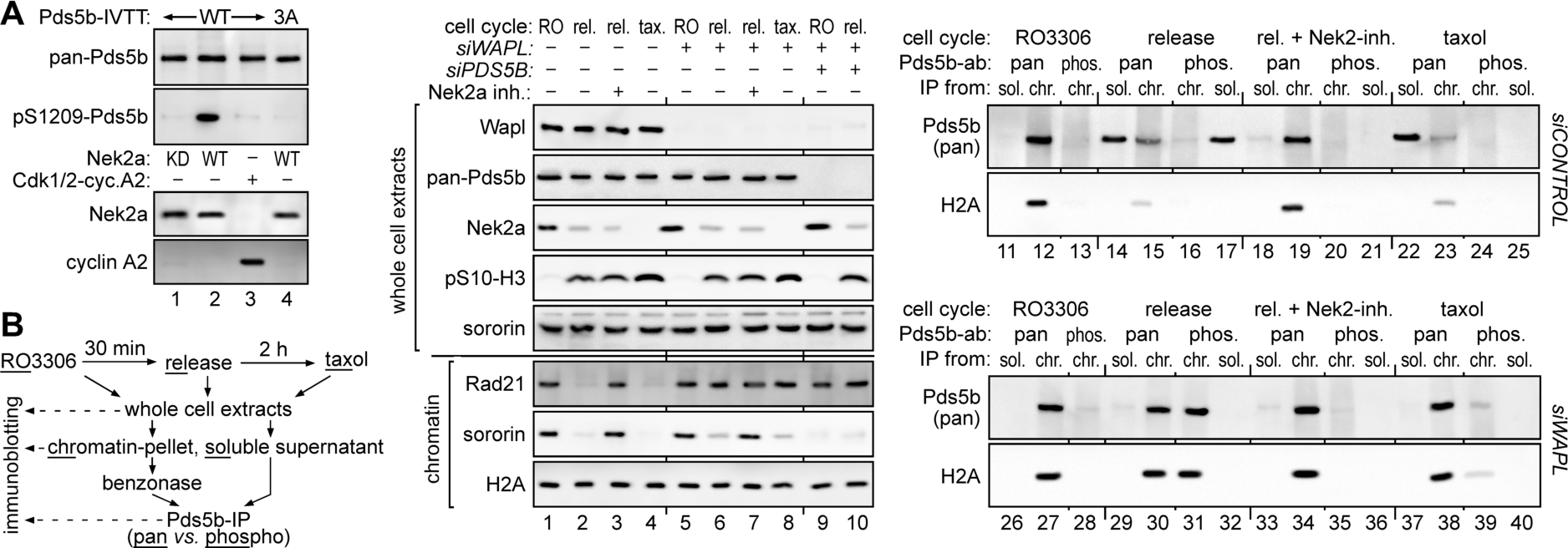
Ser1209 of Pds5b is phosphorylated by Nek2a in early mitosis. **A** Ser1209 of Pds5b is phosphorylated by Nek2a but not Cdk1/2-cyclin A2. Following incubation of *in vitro* expressed Pds5b-WT or -3A with Nek2a or Cdk1/2-cyclin A2, samples were analyzed by immunoblotting using the indicated antibodies. Nek2a-Lys37Met (KD, kinase-dead) served as a negative control. **B** Ser1209-phosphorylated Pds5b exhibits Wapl-dependent displacement from early mitotic chromatin. HeLaK cells in G2-, pro- and prometaphase were fractionated into chromatin and soluble lysate and subjected to the indicated (IP-) Western analyses. Nek2a inhibitor (NCL 00017509) was added at the time of release from RO3306 and cells were harvested 30 min thereafter.

**Figure EV4.**
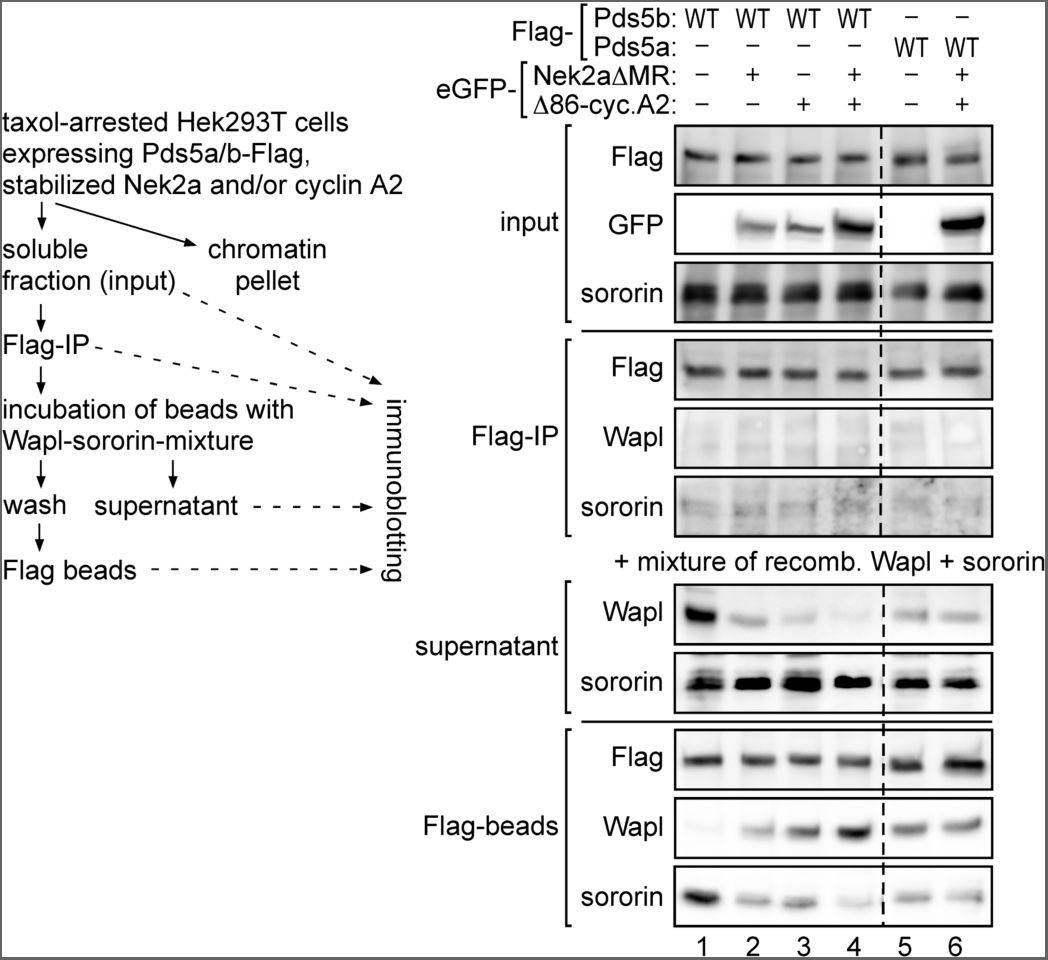
Nek2a and Cdk1/2-cyclin A2 convert Pds5b from a sororin- into an Wapl- interactor but have no effect on Pds5a’s binding behavior. Shown are the experimental outline and corresponding immunoblots. Note that eGFP- Nek2aΔMR and eGFP-Δ86-cyclin A2 migrate at the same height in SDS-PAGE.

